# Multiple Temporal and Semantic Processes During Verbal Fluency Tasks in English-Russian Bilinguals

**DOI:** 10.1101/2020.12.14.422659

**Authors:** Alan J. Lerner, Michelle Crough, Steven Lenio, Wojbor Woyczynski, Frances M. Lissemore

**Affiliations:** Neurology Department, Case Western Reserve University School of Medicine, Cleveland, OH, USA; Neurology Department, University Hospitals Cleveland Medical Center, Cleveland, OH, USA; Case Western Reserve University School of Medicine, Cleveland, OH, USA; Department of Neurology, University of Colorado Anschutz Medical Campus, Aurora, CO, USA; Department of Mathematics, Applied Mathematics and Statistics, and Center for Stochastic and Chaotic Processes in Science and Technology, Case Western Reserve University, Cleveland, OH, USA

**Keywords:** semantic fluency, bilingualism, clustering, cognition, stochastic modeling

## Abstract

Category fluency test (CFT) performance is sensitive to cognitive processes of executive control and memory storage and access, and widely used to measure cognitive performance especially in early Alzheimer’s Disease. Analytical variables have included the number of items named, and various methods to identify and quantify clusters of semantically related words and cluster switches. Also encoded in the response sequence are temporal patterns as shown by “bursts” of responses and pauses between items, that have not been received attention in determining cluster characteristics.

We studied a group of 51 adult Russian-English bilinguals and compared CFT responses based on two clustering methodologies: the semantic-based method (SEM) and a novel method based on the time interval between words (TEMP) with 8 different intercall time thresholds from 0.25 sec-15 sec. Each participant performed the task in both languages. Total number of words and cluster count was greater in Russian than English for both scoring methods, but cluster size did not differ between languages. We also studied stochastic modeling characteristics based on detrending of the “exponential exhaustion” effect seen with CFT, with most notable that total recall capacity (N∞) was greater in Russian than English (P<.05). Multiple demographic variables, and recent and lifetime usage of each language, affected both cognitive performance as measured by the Montreal Cognitive Assessment (MOCA; given in English only). Differential performance is driven by differences in demographics, more words stored in memory, and semantic and timing recall strategies.

## Introduction

Category fluency testing (CFT) is a measure of verbal fluency often employed in clinical and neuro-linguistic assessments. A subject is asked to name as many members of a category as possible, e.g. “animals”, in a given time, generally 60 seconds. The number of responses produced on the animal naming test of category fluency is widely thought to reflect an individual’s ability to produce clusters of semantically related animal names and to rapidly switch between clusters of names [1]. This ability depends on a wide range of cognitive processes including lexical access speed, executive function, education, and the size of an individual’s vocabulary [2–5]. It is also well known that verbal fluency responses per unit time are subject to “exponential exhaustion” when response numbers per unit time are binned [3]; our previous work has extended these findings by use of statistical methods to detrend the data as shown by using stochastic modeling of response times between young and old subjects, and between older adults with normal cognition and those with varying degrees of cognitive impairment [6,7]. Thus, it appears that multiple cognitive processes are working simultaneously and contribute to the simplest output measure, the total number of words recalled in 60 seconds (N_60_).

To assess clustering ability and to better understand these cognitive processes and how they contribute to semantic fluency, multiple methodologies and subject populations have been employed using semantic fluency testing [1, 8–11]. Previous studies in bilingual populations have shown small differences between languages, but have often included individuals with bilingualism of varying languages and different methods ascertaining usage between languages. Troyer et al. [1] established a method of analyzing cluster-switch data that has been widely used in studies of semantic fluency. Published studies have often relied on two raters independently assessing a sequence of responses and determining whether consecutive responses are part of the same cluster.

Historically, cluster-switch analyses have been based on the supposition that responses are generated in “bursts” with a pause before the respondent continues with another burst of responses, and that these bursts contain semantically related words. Combining “burstiness” and semantic relatedness led to the prevailing notion that related items are stored in semantic memory such that they are accessed in rapid succession. A number of studies examining the sequence and patterns of category fluency responses have shed light on how semantic memory is organized and accessed, but significant debate still exists over the utility of category fluency to study semantic structure [6] [12] [13] [14].

Here we report both semantic (SEM) and temporal clustering (TEMP) the response sequences in an animal naming task in bilingual Russian-English cognitively normal adults. Combined with the MOCA and demographics and response-related temporal variables gives a multi-dimensional view of understanding verbal fluency output as a composite measure of multiple factors.

## Methods

### Recruitment

The Institutional Review Board of University Hospitals Cleveland Medical Center approved this study; IRB ID #05-13-13, and written informed consent was obtained for all participants prior to start of study procedures.

Participants were recruited from community recreation centers and residential retirement facilities known to serve populations of foreign-born citizens in suburban Cleveland, OH. All procedures were approved by the University Hospitals Institutional Review Board prior to recruitment. Participants were interviewed individually out of hearing range from other persons in the interview area, and were compensated with a $25 gift card. Each person informed about the study was asked whether they spoke any language in addition to English. From this exchange the interviewer was able to determine whether the speaker’s English was adequate to participate, and if so then the person was consented out of hearing range of other persons. Consent process includes questioning the potential participant to assess their understanding of the study. Demographic information collected included age, place of birth, age at the time participant moved to the United States, and age when participant began learning English. Language dominance was self-reported by the participant.

### Russian and English Usage Index and Education

We attempted to model bilingualism as a continuous variable rather than a dichotomous variable, since there is no standard quantitative threshold of “bilingualism”. Participants were asked to estimate the relative use of each language by decade over their lifespan and over the past year (“recent” (English or Russian) usage). This gives a rough approximation of bilingualism as a continuous rather than a discrete variable which varies with age and life experience. We also obtained a self-report of age participant began speaking English.

Education completed was divided into five categories as follows: Less than High school graduate (1), High school graduate (2), some college (3), college graduate (4), post-graduate education (5).

### Testing Procedures

The animal naming task was administered twice to each participant, once with responses in English and once with responses in Russian. Between the two trials, participants completed the Montreal Cognitive Assessment (MOCA) in English [15], and the order of languages for the CFT (English first or Russian first) was randomized to control for priming effects by the first trial of the second.

The Montreal Cognitive Assessment (MOCA) and animal naming tests were recorded using a handheld digital device. The recordings were transcribed and the time from the start of the trial to the start of each word (elapsed time) was calculated using WavePad Sound Editor (NCH Software Inc., Greenwood, CO).

Responses from the Russian trials were translated into English by a native Russian speaker, and we recorded the total number of non-repeated responses not including errors in 60 seconds. Two raters scored each trial for semantic clustering (“SEM” method), following the Troyer et al. [1997] method with the following exceptions: we did not assign any response to more than one cluster, we counted cluster size as the number of words in a cluster, and we counted single words (i.e. those not semantically associated with a response preceding or following) as a cluster size of one. Pearson correlation coefficient (r) between raters for semantic cluster scoring was 0.9 for the English CFT, and 0.84 for the Russian CFT.

### Clustering procedures: Semantic (SEM) and Time (TEMP)

For the both scoring methods, the intercall times (time between the start of consecutive responses) was recorded. Clusters based on time (temporal clusters or “TEMP”) were analyzed without regard to the semantic relationship among responses. In developing this new approach to clustering, it is recognized that there is no standardized intercall duration threshold known to be optimal. If a duration shorter than the minimum was chosen, then each item would be its own cluster of a single word. At the far end, thresholds greater than the maximum intercall time up to 60 seconds would perforce result in a single cluster. Therefore, we analyzed the data using thresholds of 0.25, 0.5, 0.75, 1, 1.5, 2, 5 and 15 seconds.

Mean and median cluster size (number of words in a cluster) across time duration thresholds, the average cluster size (N_60_ / # clusters) were calculated for SEM scoring method, and Median TEMP cluster size in both languages.

Additionally, we determined two additional temporal variables. The initial latency is the duration from 0 seconds to first item named. We also calculated the duration from time at last item named to 60 seconds. In previous work [7] this time correlated well with total items named. Fig 1 illustrates the complex relationship of semantic versus temporal clustering for a single subject.

**Fig 1.**
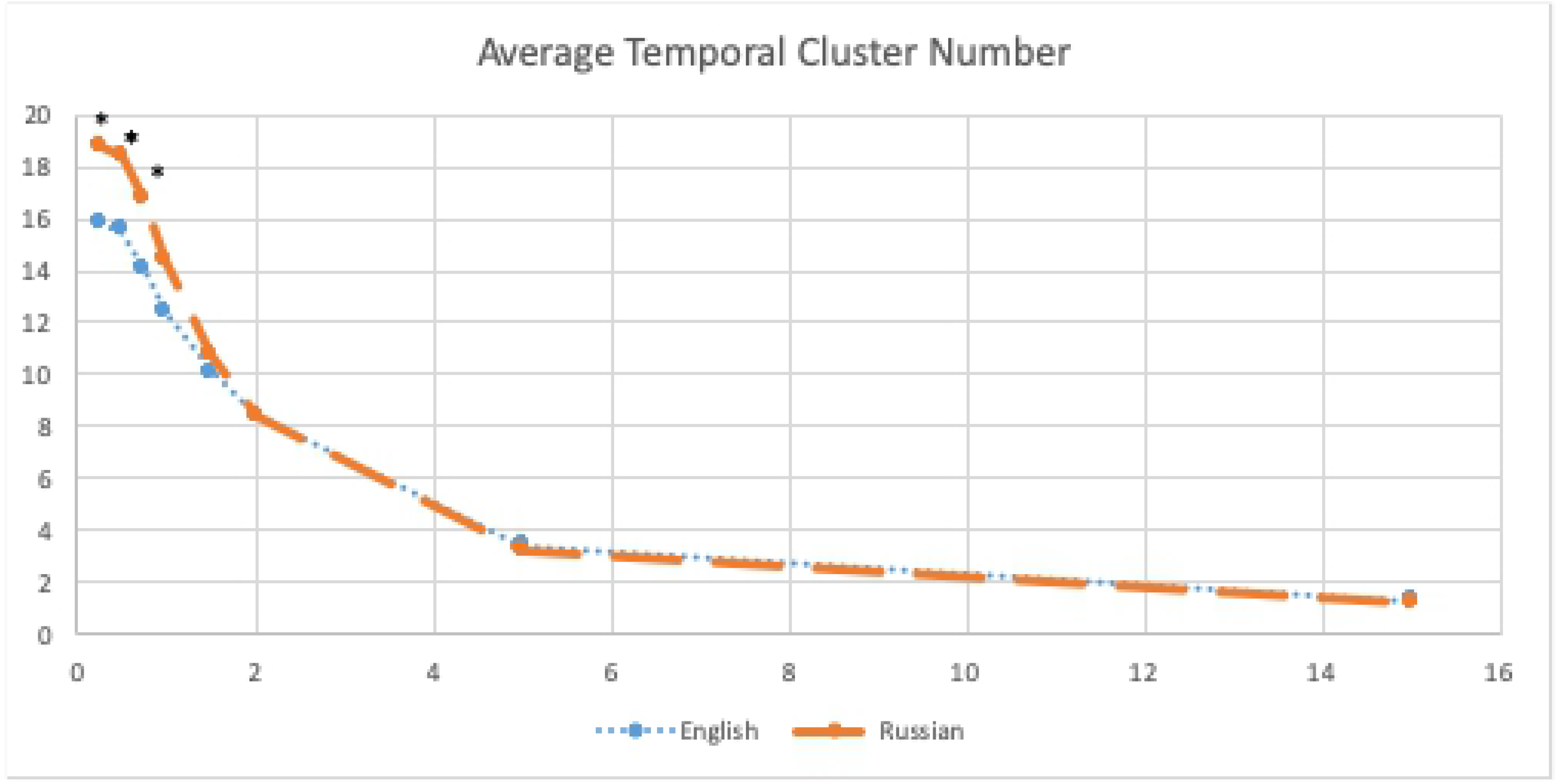
Composition of time-based cluster and semantic-based clusters in an animal naming task. The first nine responses (dog through camel) constitute the only time-based cluster in this example; those same nine responses make up three semantic-based clusters (green and yellow markers). The remaining nine responses make up two additional semantic-based clusters and five un-clustered words (gray markers).

### Temporal Detrending Variables and Statistical Analysis

Participant responses were analyzed using detrending procedures as described by Meyer et al, 2012. This creates derived variables N_60_ (the number of words recalled), N∞ representing individual’s “total recall capacity” allowing infinite time for recall, or the rate at which the subjects responses approach an asymptote; We must emphasize that the parameter N∞ is called here the “total recall capacity” only figuratively, with quotation marks applied advisedly. The actual recall process cannot possibly extend its exponential behavior to infinite time as a matter of both mathematics and common sense. Accepting the unlimited exponential behavior would practically mean that after, say, one hour the individual’s recall ability would be essentially zero, an obvious nonsense. So N∞ is just a useful parameter in the exponential exhaustion model. Tau (τ), which is the exponential time “latency” constant. The latter can be conveniently thought of as the time by which the individual reaches e^−1^ = 36.8% of their “total recall capacity”.

Additionally, the distribution of the detrended intercall times approximates the Weibull stretched exponential distribution and the three parameters of the distribution were calculated for each participant with sufficient responses [6]. The three components of the Weibull distribution: gamma, which is related to speed of response; beta which relates to the shape of the response distribution and eta, a scaling factor. Statistical analysis utilizing summary statistics, one-way ANOVA, univariate Spearman correlations and non-parametric statistics were done using JMP 14.0.

## Results

### Demographics

Table 1 shows the demographic characteristics of the subject cohort. More than 70% of participants were female, ranged in age from 19 to 75 years, were well educated (86% graduated college or had a post-graduate education), and all but two were born outside the United States, primarily in Russia or Ukraine. All participants spoke Russian before they spoke English, and began English language instruction between the ages of 3 and 59 years. Table 1 shows subject demographics, MOCA scores, MOCA letter fluency word count and N_60_ in each language. Table 1 also shows the life time and previous year index of usage of each language. Table 2 shows the univariate correlation analysis showed that both Russian and English word counts (N_60_) correlated significantly between themselves, and were highly correlated with MOCA score and MOCA letter fluency, education, lifetime Russian shown).

**Table 1:**
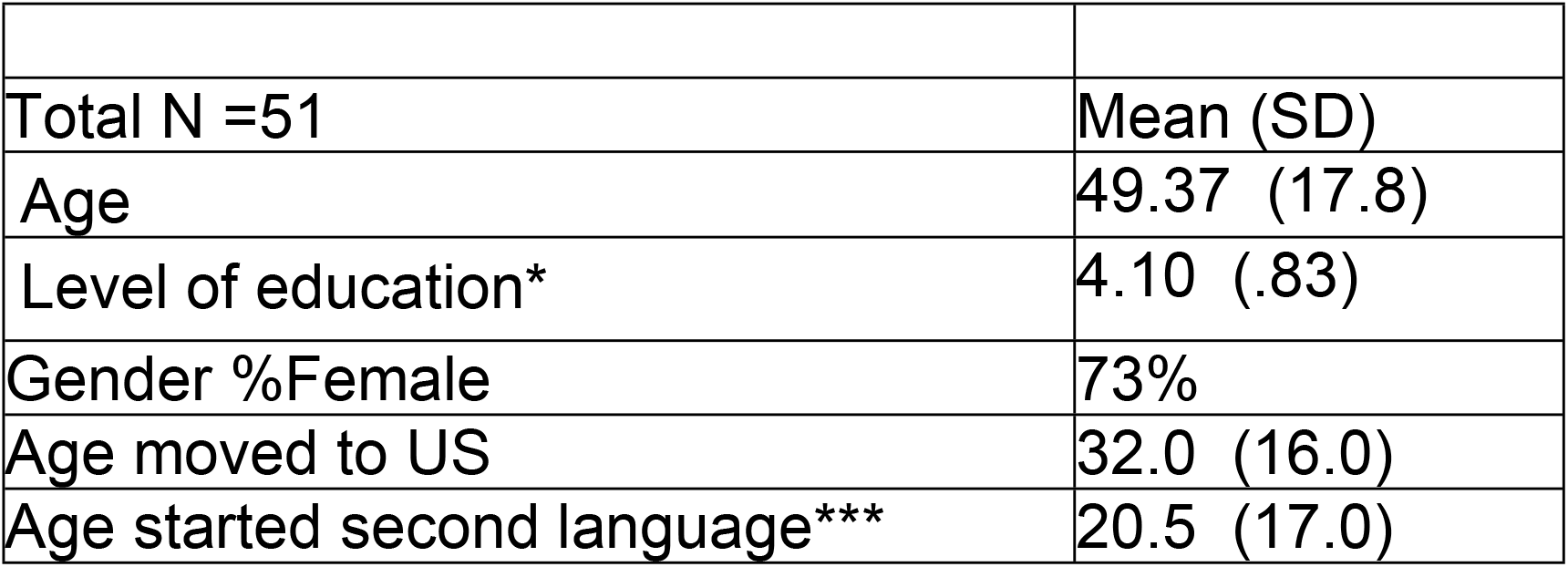
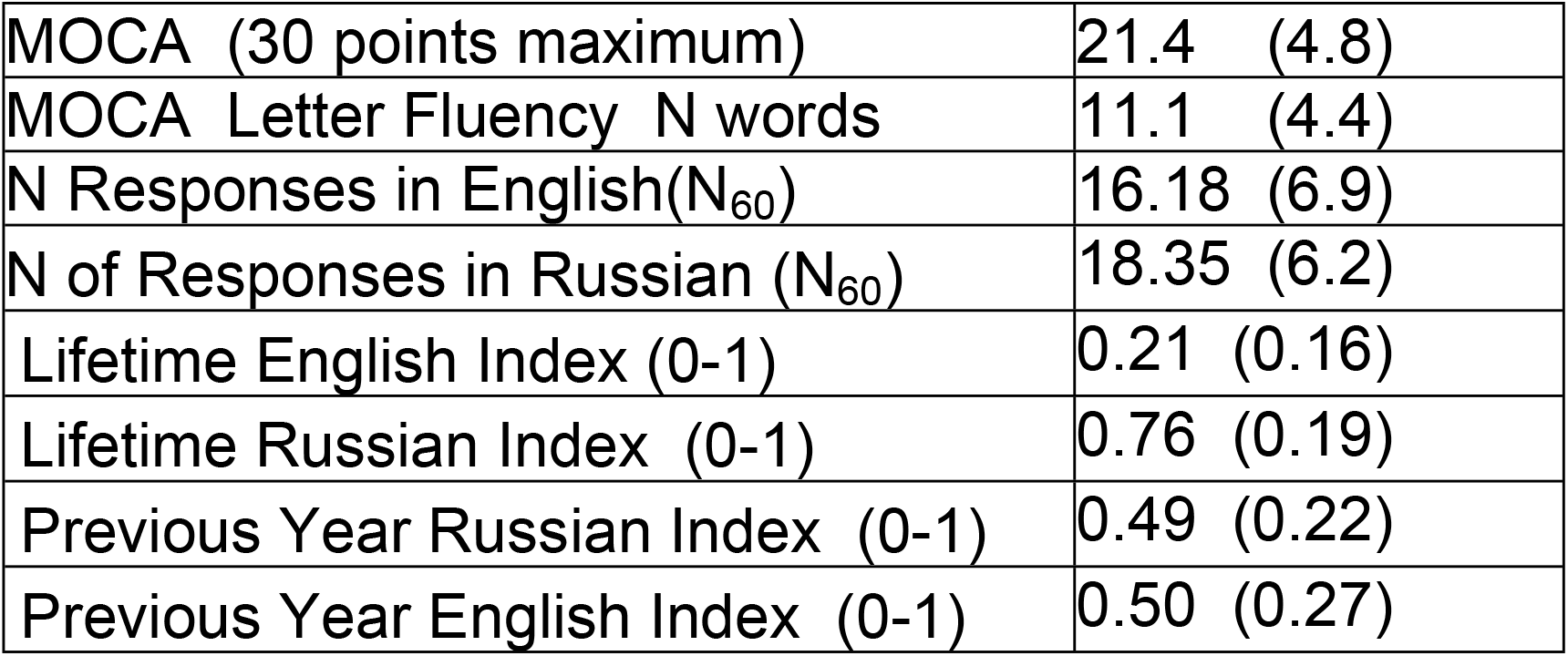
Subject Demographics.

**Table 2:** Univariate correlations (Spearman’s rho) of word production in each language and demographic variables and the Montreal Cognitive Assessment scores.

|  | N60 English | Age | MOCA Score | MOCA Letter Fluency | Lifetime Russian Index | Most Recent Russian Index |
| --- | --- | --- | --- | --- | --- | --- |
| N<sub>60</sub> Russian | 0.43** | -0.14 | 0.37** | 0.41** | -0.24 | 0.08 |
| N<sub>60</sub> English |  | -0.61*** | 0.73*** | 0.53*** | -0.54** | -0.26 |
| Age |  |  | -0.62*** | -0.17 | 0.50** | 0.44* |
| MOCA Total Score |  |  |  | 0.59*** | -0.48** | -0.24 |
| MOCA Letter Fluency Count |  |  |  |  | -0.28 | 0.68* |
| Lifetime Russian Index |  |  |  |  |  | 0.81*** |

NS=Not significant; \*<0.05, \*\*<0.01, \*\*\*, <001

**Table 3.** N_60_ characteristics and the lack of priming effects based on first language tested for verbal fluency output.

|  |  | N | N<sub>60</sub> mean(sd) |  |
| --- | --- | --- | --- | --- |
| Language | Russian | 51 | 18.4(6.2) | t=16.75,<br>p<.0001 |
|  | English | 51 | 16.1 (6.9) |  |
| Language x Trial |  |  |  | F,p |
| Russian Responses | First Trial Russian | 24 | 17.8 (6.1) | 0.31, NS |
|  | First Trial English | 27 | 18.8 (6.4) |  |
| English Responses | First Trial Russian | 27 | 16.5(7.7) | 0.12, NS |
|  | First Trial English | 24 | 15.9 (6.2) |  |

### Comparing methods visually

Fig 1 displays the outcomes of the two clustering methods for one participant’s Russian responses in the CFT. The difference in the pattern of clusters between the two methods is striking, especially in the first 9 responses, which are grouped as one TEMP cluster and three SEM clusters. The remaining graphs for all participants’ Russian responses and English responses are shown in S1 Fig and S2 Fig.

### Response characteristics

There was no difference in the number of words produced in English or Russian tested first versus English or Russian tested second in the CFT trials, indicating there was no priming effect of repeating the test within a short time. Likewise, the order of languages in the two trials (Russian first or English first) did not make a difference in the number of words produced in that language.

Table 4 shows the comparison between languages of the detrended response variables. N∞ was significantly larger in Russian, suggesting that the pool of available responses was larger, and thus one factor for greater number of word responses in Russian (see table 1)

**Table 4.**
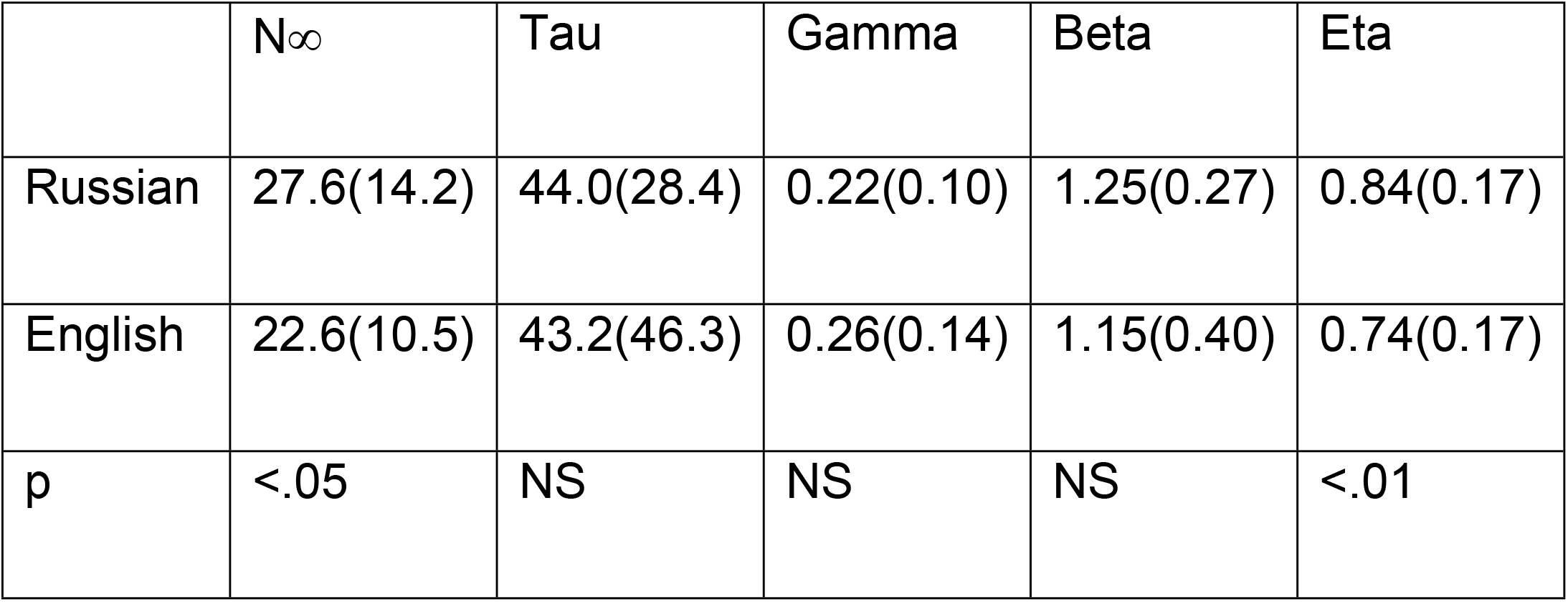
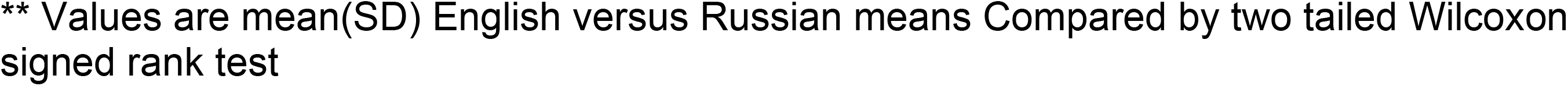
Temporal Recall Indices in Russian and English (N=48).

|  | N<sub>∞</sub> | Tau | Gamma | Beta | Eta |
| --- | --- | --- | --- | --- | --- |
| Russian | 27.6(14.2) | 44.0(28.4) | 0.22(0.10) | 1.25(0.27) | 0.84(0.17) |
| English | 22.6(10.5) | 43.2(46.3) | 0.26(0.14) | 1.15(0.40) | 0.74(0.17) |
| p | <.05 | NS | NS | NS | <.01 |

\*\* Values are mean(SD) English versus Russian means Compared by two tailed Wilcoxon signed rank test

Table 5 shows the univariate correlations (Spearman’s rho) between word count, N∞, tau, average number of clusters for each clustering method, average cluster size (SEM) and median cluster size (TEMP). N∞ correlated best with word count and clustering variables and Tau, but the correlation with temporal cluster number was not significant in Russian, while it was in English testing. Interestingly, most other variables except word count in Russian median N TEMP clusters were not significant.

**Table 5:** Univariate correlations of word count, clustering variables and detrended time variables.

| | | N<br>SEM<br>clusters | N Median<br>TEMP<br>clusters | Mean<br>SEM<br>Cluster<br>size | Median<br>TEMP<br>cluster<br>size | $N_{\infty}$ | Tau |
| --- | --- | --- | --- | --- | --- | --- | --- |
| Russian | Word count ($N_{60}$) | 0.78*** | 0.75*** | 0.26 | 0.56*** | 0.74*** | 0.36* |
|  | N Semantic Clusters |  | 0.53** | -0.32* | 0.44** | 0.52** | 0.20 |
|  | Median N TEMP clusters |  |  | 0.14 | -0.02 | 0.67*** | 0.51** |
|  | Mean SEM Cluster size |  |  |  | 0.13 | 0.38** | 0.27 |
|  | Median TEMP Cluster size |  |  |  |  | 0.26 | -0.07 |
| | $N_{\infty}$ | | | | | | 0.81*** |
| English | Word count | 0.72*** | 0.83*** | 0.44*** | 0.64*** | 0.81*** | 0.04 |
|  | N Semantic Clusters |  | 0.68*** | -0.24 | 0.31 | 0.55*** | 0.12 |
|  | Median N TEMP clusters |  |  | 0.22 | 0.16 | 0.63*** | 0.14 |
|  | Mean SEM Cluster size |  |  |  | 0.44** | 0.43** | -0.01 |
|  | Median TEMP Cluster size |  |  |  |  | 0.47** | -0.15 |
| | $N_{\infty}$ | | | | | | 0.45** |

$p < .05$ ; \* $p < .01$ ; \*\*\* $p < .001$

**Table 6.** Clustering Characteristics by Language and Methodology of Clustering (Semantic versus Temporal).

|  | N<br>Semantic<br>Clusters | Semantic<br>Mean<br>Cluster<br>size<br>(N<br>words) | Temporal Clusters (Mean (SD)) |  |  |  |
| --- | --- | --- | --- | --- | --- | --- |
|  |  |  | N Mean | Mean<br>Cluster size | N Median | Median Cluster size |
| Russian | 9.63<br>(3.4) | 2.09<br>(0.6) | 11.5 (2.8) | 1.62 (0.18) | 12.6 (3.0) | 1.49 (0.30) |
| English | 8.1(3.3) | 2.04<br>(0.7) | 10.125<br>(3.3) | 1.53 (0.20) | 11.2 (3.6) | 1.39 (0.29) |
| p-value | <.0001 | NS | <.01 | <.02 | <.02 | NS |

### Comparing Cluster Characteristics

When comparing cluster characteristics between languages, semantic cluster count was significantly higher in Russian than in English (Table 5). This is likely an effect of the greater N_60_ in Russian than English; more words generally result in more clusters. Cluster sizes, however, did not differ significantly between Russian and English (Table 5). There were significantly more temporal clusters in Russian than English for all durations 1 second or less. However, the curves were of similar shape, and 0.25 sec threshold duration had cluster counts that approximated N_60_, and threshold duration of 15 seconds almost always yielded a single cluster of all words (Fig 2).

**Fig 2:**
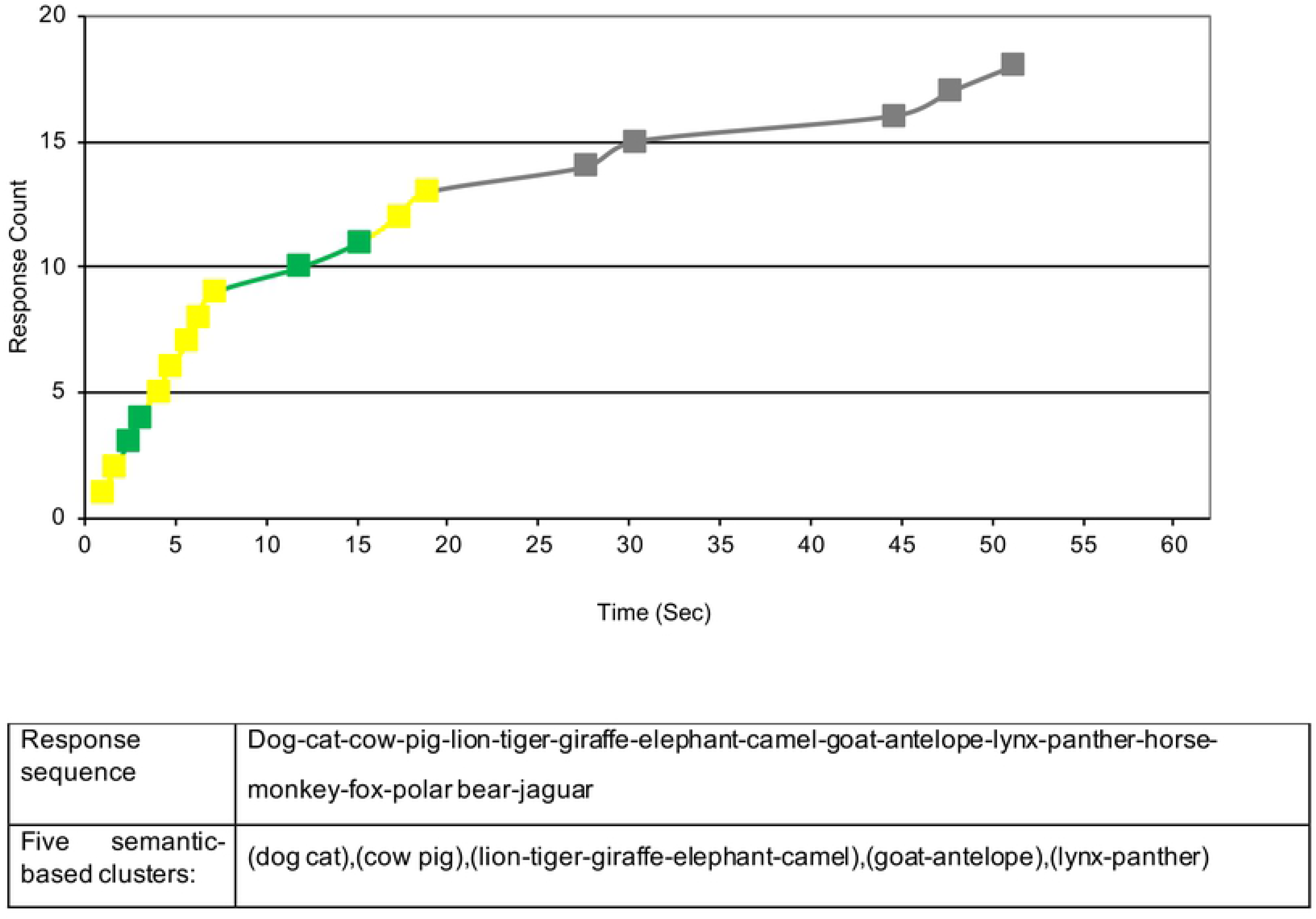
Average Number of Clusters based of Temporal Duration Thresholds by Language.

### Initial latency and terminal duration

The initial latency was significantly faster in Russian than English (1.14 ± 0.58 vs. 1.94 ± 1.70 seconds; p<0.003). Time from last word to 60 seconds was significantly less in Russian than English (6.85 ±6.1 sec vs. 9.62 ± 8.6 sec; p<.05). Both of these differences probably contribute to the greater number of words and ultimately clusters in Russian than English. Several participants had terminal durations of more than 15 seconds, and two participants had terminal durations in English of more than 30 seconds. They are also consistent with the smaller gamma response variable in Russian than English, although that difference was not statistically significant.

## Discussion

Given the many analytical methods applied to CFT in the literature, it is clear that an enormous amount of information is encoded in the item content, as well as timing intervals between words. The major focus and results of this study involve comparison of the CFT in a cohort of Russian-English Bilinguals tested in both languages, and comparison of temporal and semantic cluster scoring methods using different threshold durations for defining temporal clustering. Since both semantic and temporal information are simultaneously encoded in the response sequence, it is important to determine their relationships on a quantitative level. Our study’s major findings relate to the two main aims of the study: comparison of semantic and temporal processing between languages, and the feasibility of measuring temporal clustering. For the former aim, participants produced more responses in Russian than English, and this appears multiply determined, including demographics and differential language use, but also differences in response timing, total time spent engaged in task, and size of lexicon in each language as measured by total recall capacity. For the latter aim, measuring temporal clustering is quite feasible and allows comparison to the established semantic clustering method.

Temporal clustering can only be done and measured when the CFT is recorded continuously, rather than the binning method used in many previous studies [3], [16]. Use of different duration thresholds to define temporal clusters showed a similar pattern of temporal clusters in both languages, and a maximum number of temporal clusters occur using a 1.5-2 sec cutoff of the intercall duration to separate clusters.

Previous literature has expressed concern about the subjective aspects of determining semantic clustering, although the many studies using variants of Troyer’s methodology have shown differences consistent with the known neurobiology of neurodegenerative disorders such as AD [9] [17]. These concerns arise from the ambiguity involved in determining semantic relatedness. Thus, a sequence of *dog-cat-parrot-fish-whale* could be interpreted as two clusters (dog-cat-parrot (Pets); fish-whale (Marine animals), or perhaps dog-cat-parrot-fish (Pets) and whale (marine mammal), or as three clusters of dog-cat (pets), parrot (bird) and fish-whale (marine animals).

Long duration pauses in the response sequence are common, and there is often a “second wind” phenomenon, with a second acceleration of responses after a long pause - in effect, restarting the task. These longer duration pauses are problematic since they suggest alterations in brain processing whose meaning is ambiguous. In the SEM method but not in the TEMP method, a long pause is incorporated into the sequence of a cluster raising the question of whether the respondent “intended” the responses to be semantically related. That is, long durations finally producing a semantically related word, may indicate the end of one cluster, and then essentially restarting the semantic association process anew. The TEMP method more accurately reflects how respondents verbalized their responses and allows the pauses themselves to be utilized in analyses of the retrieval process instead of “concealing” pauses within clusters as in SEM. Another point related to this is that long duration pauses or stopping the response sequence early effectively turn the 60 second test into a much shorter test and heavily affect the number of items produced, which is the simplest analytical method for evaluating semantic processing. Also to be considered in language function is the initial latency of the first word and the duration from last word to 60 second end of test. Particularly the latter contributes a constraint to increasing the number of words produced, effectively shortening the 60 second test, occasionally by as much as 30 seconds. Whether this is a motivational or attentional or linguistic issue cannot be determined from the available data. The lower initial latency in Russian probably was one factor contributing to increased word production in that language.

Hills, et al. 2015 [18] proposed two alternative but not mutually exclusive models for semantic memory search. Their associative model is based on “a connected sequence of related items”, presumably connected by frequency of usage even if the items are not closely related semantically. Second, their categorical model relies on recalling “entire predefined categories” and choosing responses from within that group. Our data may support their associative model. Sequential responses often show little semantic relatedness, for example in Fig 1 where “pig” is followed by “lion” (same TEMP cluster, different SEM cluster). Hills et al. [18] refer to “low-similarity transitions”; our data show transitions between responses that are short in time but do not necessarily have “short” semantic connections. That observation supports the idea that TEMP clusters correspond to the associative model because high-usage responses are likely to show up in the same cluster even if they are not closely related semantically. Hills et al. conclude that retrieval from semantic memory is a process both of frequency of usage and of categorical similarity, and long pauses between semantically related responses (as seen in our data) support that idea. Other forms of semantic clustering using different word retrieval constructs have been compared by Abenwender, Swan, Bowerman and Connolly [19] but further discussion is beyond the scope of this study.

Another methodological concern of both methods involves the treatment of single word “clusters”. By definition, the first word produced is part of a cluster, and if the subsequent response exceeded the temporal threshold, a cluster of a single word is generated. Thus, even the definition of “temporal cluster” contains some ambiguity. Whereas semantic clusters are derived from two independent raters, definition of temporal clustering is done automatically, but cluster numbers vary depending on the threshold duration chosen.

Troyer et al 1997 [1] used number of switches between clusters as a proxy for direct cluster count, stating “(s)witches were calculated as the number of transitions between clusters, including single words….”; i.e. single word clusters. Dramatic differences in count caused by cluster definition has consequences in neuro-linguistic assessments insofar as cluster count is considered a reflection of cognitive function. Haugrud et al. [9] note “small changes in scoring…can change…measures of clustering, (hence) average cluster size might not be the most effective method for differentiating AD from healthy aging” [20]. A one-word cluster has no semantic association with a word preceding or following it, and it often exceeds in time an association with adjacent responses. Therefore, it is also possible that single word clusters do not totally fit SEM or TEMP criteria, and that analyses of clusters>1 word alone may point towards more meaningful conclusions regarding semantic memory structure and access.

The cultural, educational and life experiences of the respondent may influence how frequently words are used and how readily they are retrieved from memory, and similarly, those of the rater may influence cluster composition. Interestingly, we found a lower correlation between our raters (U.S.-born native English speakers) in semantic category scoring in Russian (after translation) (Pearson r=.84) than in English (Pearson r=.90). This suggests it was more difficult for raters to consistently infer the response relationships intended by respondents in the language not spoken by our raters.

Temporal clustering methods may have a theoretical advantage over semantic methods in terms of understanding neural function as reflected by advanced statistical methods. The process of semantic recall for related terms is similar to how animals search for food in resource patches (for example bees in a flower patch), which is the focus of optimal foraging theory (OFT) [21]. Optimal foraging models posit that animals search patches such that foraging efficiency is maximized. Likewise, “patches” of semantic memory are searched for unique animal names in the CFT. OFT has recently been applied to analyses of response sequences in CFTs [18], divergent thinking processes [22], and to predict intercall times [23]. In addition to optimal foraging, other kinds of models have been utilized to further understand memory association and retrieval, including mathematical, physical, and computer-based analyses [24] [25] [26] [27].

Limitations of the present study include the absence of monolingual control groups in each language, and of a general vocabulary assessment in either language before testing. Our study population, however, was highly educated (Table1) which likely indicates strong vocabulary ability. The sample of participants is a convenience sample, and may not reflect the wide population of Russian-English or other bilingual combinations. Application to aging and disease models also awaits further study.

Our prior work examined CFT in groups with varying levels of cognitive impairment [7] focused on intracluster and intercluster timings but did not include analysis of temporal clustering. In that study, cluster size did not vary significantly across groups, but cluster counts did, a pattern similar to what we found in the current analysis. Clusters with more words means fewer clusters (i.e. fewer cluster switches), and cluster switching has been used as a proxy for executive function [8] [28] [29] [5]. We could likely expect that other types of category fluency, e.g. vegetables, four legged animals, food or clothing, could be analyzed by the TEMP method as well.

Category fluency testing is a clinically useful measure because of the enormous amount of encoded information utilizing multiple brain processes contributing to its outcome. Integrative tests, such as gait timing, clock drawing or CFT are useful screening tools precisely because performance integrity implies intact brain processing, and conversely, it is sensitive to many types of baseline neurological ability, brain injury and cognitive decline beyond Alzheimer’s disease and related disorders [30] [31]. As reviewed here, clustering reflects multiple brain processes, and both semantic and temporal clustering provide insights into these very brain processes. Temporally-based cluster scoring method for the animal naming task is equally feasible and possibly less ambiguous than the semantic-based method although the optimal threshold duration between items, used to define “clusters” is varies between respondents. Additionally, temporal clustering may reduce basic inter-rater reliability because the start of a cluster and its end are quantitatively determined, obviating semantic relatedness judgements. This method also allows for a faster scoring process which can be easily adapted to automated programming. Future studies in different populations are needed to define the relative contributions of the two methods in determining the clinical and research significance of each.

## Supporting Information

**S1 Fig. Semantic and temporal cluster compositions for responses in Russian in the animal naming task.** Response count is graphed over time (60 seconds), and accompanying chart shows assignment of semantic clusters and of temporal clusters. The top line of the chart identifies participant ID number and response count, with sequence of animal names in the second line. Cluster switches are indicated by alternate shading of boxes below the response sequence (semantic clusters-- third line; temporal clusters-- fourth line); no shading under a word indicates it was not part of a cluster.

**S2 Fig. Semantic and temporal cluster compositions for responses in English in the animal naming task.** Response count is graphed over time (60 seconds), and accompanying chart shows assignment of semantic clusters and of temporal clusters. The top line of the chart identifies participant ID number and response count, with sequence of animal names in the second line. Cluster switches are indicated by alternate shading of boxes below the response sequence (semantic clusters-- third line; temporal clusters-- fourth line); no shading under a word indicates it was not part of a cluster.

## Acknowledgements

This study was funded by the Brain Health and Memory Center, Neurological Institute of University Hospitals Cleveland Medical Center. The authors wish to thank S. Klayman for translating responses from Russian.

